# Independently paced calcium oscillations in progenitor and differentiated cells in an *ex vivo* epithelial organ

**DOI:** 10.1101/2021.08.11.454108

**Authors:** Anna A. Kim, Amanda Nguyen, Marco Marchetti, Denise J. Montell, Beth. L. Pruitt, Lucy Erin O’Brien

## Abstract

Cytosolic calcium is a highly dynamic, tightly regulated, and broadly conserved cellular signal. Calcium dynamics have been studied widely in cellular monocultures, yet *in vivo* most organs comprise heterogeneous populations of stem and differentiated cells. We examined calcium dynamics in each cell type of the adult *Drosophila* intestine, a self-renewing epithelial organ where multipotent stem cells give rise to mature absorptive enterocytes and secretory enteroendocrine cells. Here we perform live imaging of whole organs *ex vivo*, and we employ orthogonal expression of red and green calcium sensors to determine whether calcium oscillations between different cell types are coupled. We show that stem cell daughters adopt strikingly distinct patterns of calcium oscillations when they acquire their terminal fates: Enteroendocrine cells exhibit single-cell calcium oscillations, while enterocytes exhibit rhythmic, long-range calcium waves. These multicellular waves do not propagate through progenitor cells (stem cells and enteroblasts), whose oscillation frequency is approximately half that of enteroendocrine cells. Organ-scale inhibition of gap junctions eliminates calcium oscillations in all three cell types, even, intriguingly, in progenitor and enteroendocrine cells that are surrounded only by enterocytes. Our findings establish that cells adopt fate-specific modes of calcium dynamics as they terminally differentiate and reveal that the oscillatory dynamics of different cell types in a single, coherent epithelium are paced independently.

## INTRODUCTION

Calcium is a versatile signaling molecule that regulates critical cellular functions such as contraction and cellular excitability in all organ systems ^1–3^. Changes in calcium signal dynamics have been linked to crucial cell behaviors, among them intercellular communication, cell cycle and proliferation, and migration ^4–7^. Intracellular calcium concentrations can also regulate cellular responses and physiology by modulating signal transduction pathways such as MAPK^8,9^ and IP3^10–12^. In excitable tissues, such as in electrically coupled heart muscle cells, calcium plays a central role in impulse propagation for coordinated pacing of contractions ^13,14^. Large-scale calcium waves have been observed in cultured and in mouse hippocampus astrocyte networks ^15,16^. Studies in *Drosophila* demonstrated intercellular calcium waves that traverse large tissue domains and may depend on actomyosin organization^17^. These tissue-level calcium dynamics, which occur in imaginal discs––the tightly coupled epithelial structures that give rise to the external structures of the adult fly, were implicated in organ growth and size modulation ^18,19^. Furthermore, intercellular calcium waves were shown to be induced mechanically in this developing epithelia ^20^.

While calcium dynamics have been widely studied in cellular monoculture, investigations in tissues composed of multiple cell types have been limited, particularly for non-excitable tissues. Recently, blood progenitors in the *Drosophila* lymph gland were shown to form a gap junction-mediated network that can regulate calcium signaling ^21^. Here we establish the adult *Drosophila* intestine as an *ex vivo* model for studying diverse calcium dynamics that occur simultaneously in a mature organ. In the adult fruit fly midgut, calcium signaling is an important regulator of stem cell activity ^22,23^. Calcium transients are mechanically induced via the mechanosensitive ion channel Piezo in a subpopulation of stem cells in the fly gut ^23^. The midgut, like most mature organs, undergoes continuous turnover in which tissue-specific stem cell divisions produce progeny that differentiate into multiple cell types. Yet how calcium dynamics regulate, and are regulated by, cell differentiation and organ-scale inputs remains largely unknown.

Here, we simultaneously express spectrally distinguishable calcium indicators in each midgut cell type and perform real-time analysis of single- and multi-cell oscillations to produce a fate-resolved, tissue-scale overview of calcium dynamics. We investigate fate-specific changes as cells differentiate in their native tissue environment. We describe rhythmic multicellular calcium waves in enterocytes and cell-type specific calcium oscillations. These results demonstrate that as cells differentiate from stem-like into distinct terminal fates, they adopt cell type-specific calcium oscillations and waves that are a hallmark of the mature organ.

## RESULTS

### All major cell types in the adult middle midgut exhibit calcium oscillations under whole organ culture *ex vivo*

To examine calcium dynamics in the adult fruit fly midgut, we performed live imaging of whole, intact organs employing the genetically encoded calcium sensors, GCaMP6s ^24^ and jRCaMP1b ^25^. To examine calcium oscillations in each midgut cell type (Figure 1A), we expressed a genetically encoded calcium sensor using the GAL4/UAS system^26^ under the control of cell type-specific drivers. We dissected midguts from mated female fruit flies aged 4-7 days and mounted them by adapting an *ex vivo* organ culture protocol (see Methods). To characterize cell-type specific calcium activity in terms of temporal dynamics, we traced mean fluorescence intensity of individual cells as a function of time (see Methods) and quantified the frequency of oscillations.

**Figure 1.**
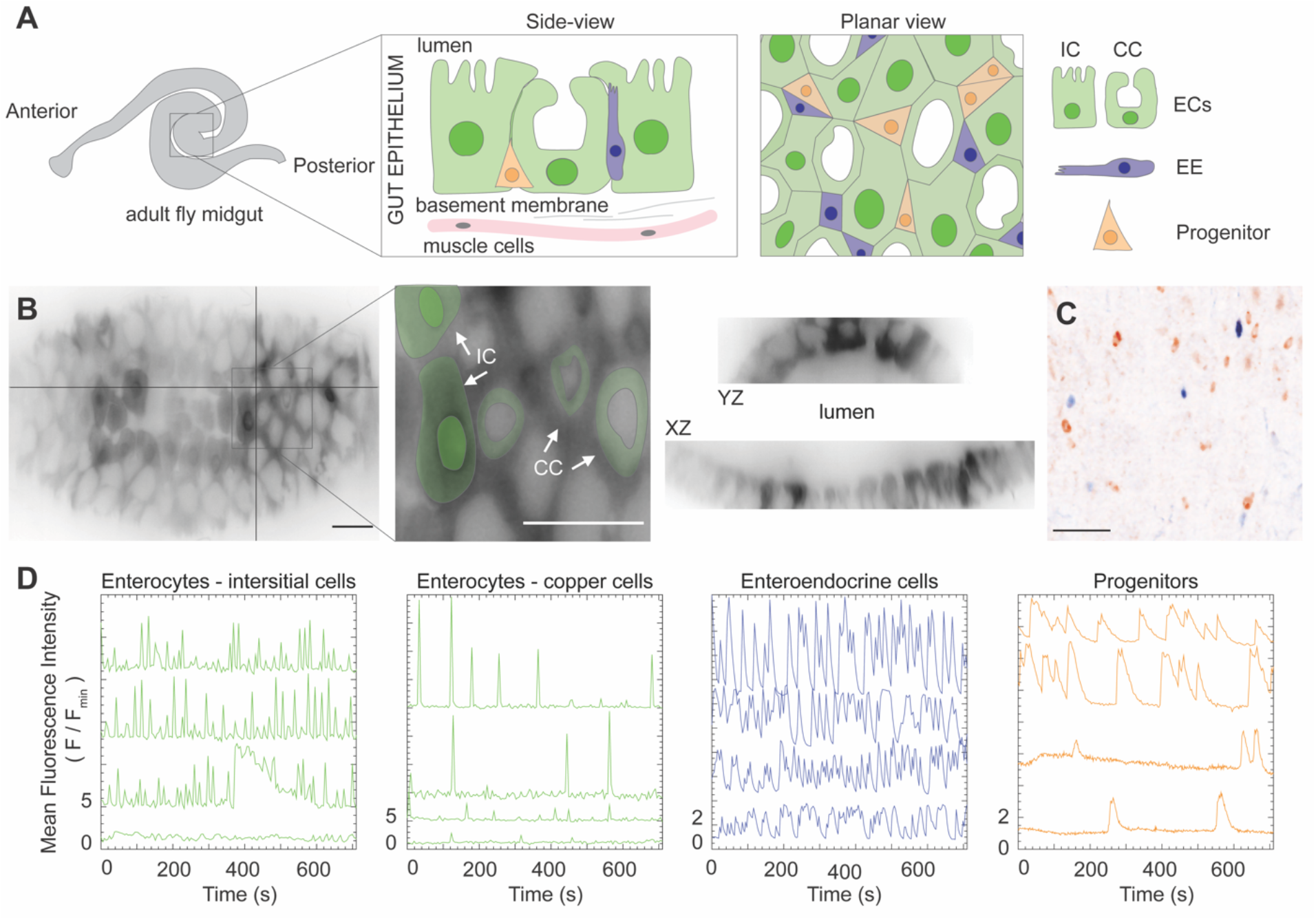
Distinct calcium oscillations in the midgut’s major cell types - enterocytes, enteroendocrine cells, and progenitors in the middle midgut region. A) Schematic of the middle (R3) midgut, the copper cell region, with side and planar views, IC: interstitial cell, CC: copper cell, EC: enterocyte, and EE: enteroendocrine cell. B) Max intensity projection and orthogonal views of a z-stack acquired with a light-sheet microscope (*mex-GAL4>UAS-GCaMP6s*). Interstitial and copper cells were distinguished by the unique shape of the copper cells, examples are marked with false-colored overlays and arrows in the inset. C) Maximum intensity projection demonstrating the distribution of enteroendocrine cells (blue) and progenitors (orange) (*esg-LexA>LexAop-jRCaMP1b, pros-GAL4>UAS-GCaMP6s*) using an inverted colormap^28^. All scale bars, 25 µm. D) Stem cell daughters acquire distinct patterns of calcium oscillations. Representative traces of fluorescence intensity of genetically-encoded calcium indicators in enterocytes (*mex-GAL4>UAS-GCaMP6s*, Movie S1), enteroendocrine cells (*prosGAL4>UAS-GCaMP6s*, Movie S2), and progenitors (*esg-GAL4>UAS-jRCaMP1b*, Movie S3). Representative traces were selected from five movies for ECs, five movies for EEs, and four movies for progenitors.

### Terminally differentiated cells of the R3 midgut exhibit distinct calcium dynamics

The midgut’s R3 region, which is responsible for acid secretion, attracted our attention due to the consistent appearance of rapid, multi-cellular calcium waves that travel through R3 enterocytes. We also observed calcium dynamics in enterocytes in the posterior region of the midgut, but not consistently. We did not observe any calcium dynamics in the anterior region. We used the GAL4 driver *midgut expression 1* (*mexGAL4)* to label enterocytes ^27^ and focused the remainder of our experiments on R3.

As a starting point, we measured the frequency of calcium oscillations in single cells of each cell type over time. Enterocytes in the R3 region are subdivided into acid-secreting copper cells (CCs) and interstitial cells (ICs) ^29–31^. For analysis, we distinguished the two sub-cell types by the copper cells’ unique shape (Figure 1B). Interstitial cells exhibited wave-like calcium dynamics with a mean oscillation frequency of 32 mHz ± 2.5 (SEM), averaged across cells. The mean oscillation frequency per gut for interstitial cells was 31 mHz ± 8.9 (SEM). By comparison, copper cells primarily displayed calcium spikes at a frequency of 4.9 mHz ± 0.7, averaged across cells (Figure 1D). The mean oscillation frequency per gut for copper cells was 5.2 ± 0.9. Interestingly, we also observed relatively rapid oscillations in fluorescence signal in enteroendocrine cells (*prospero* (*pros*) expressing) of 13 mHz ± 1.7, at a rate ∼2 times higher in comparison to progenitors (stem cells and enteroblasts, which both express *escargot* (*esg*)), 5.8 mHz ± 0.6. The mean oscillation frequency per gut was 15 ± 4.3 and 7.1 ± 2.6 for enteroendocrine cells and progenitors, respectively. Finally, the robust calcium oscillations we observed in progenitors are consistent with prior reports ^22,23^.

### Long-range calcium waves travel across enterocytes

Unexpectedly, we found long-range calcium waves that propagate across large fields of enterocytes in the R3 region of the midgut (Figure 2A). We observed high GCaMP signal shifted rapidly across several cell lengths in multiple directions (Movie S1), with the longest wave we observed covering approximately 5 cells. The calcium waves exhibited a multitude of dynamic patterns, including propagating along a single trajectory, splitting, and colliding, and they did not appear to have a clear point-of origin. Multi-enterocyte waves travel almost entirely through interstitial cells and rarely through copper cells. Despite propagating in multiple directions, the GCaMP signal recurred continuously in the same cell (Figure 2B). In this case, for example, the cells spiked on average every 23 s, leading to a frequency of 44 mHz ± 1.4.

**Figure 2.**
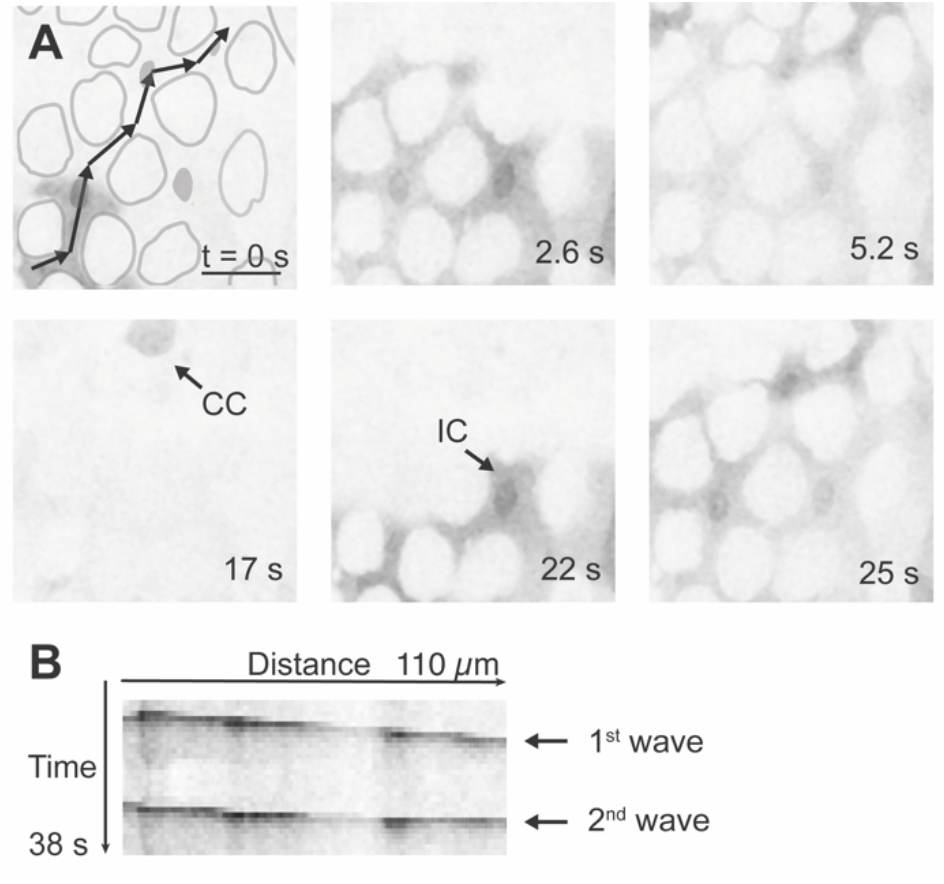
Calcium wave propagation in enterocytes. A) Propagation of calcium waves across several cell lengths (*mex-GAL4>UAS-GCaMP6s*) in a single plane. Approximations of copper cell outlines are depicted in gray. Single black arrows identify examples of copper cells (CC) and interstitial cells (IC) based on the unique shape of copper cells. The trajectory of a calcium wave (identified visually) is depicted by the linked black arrows in A. Scale bar, 25 µm. See Movie S1. B) Kymograph along the direction of the calcium wave as identified in A, demonstrating that a second wave appears and propagates in a similar direction (arrows).

To assess whether individual waves travel along recurring paths, we followed GCaMP signal in selected regions of a single midgut (Figures 3A, B and C, D), which encompassed 8-13 cells in a ∼3000 µm^2^ area. Using fluorescence intensity, we estimated cell outlines and identified copper cells based on their unique cell shape and signal (Figure 1B); interstitial cells were identified as space between the copper cells ^31,34^. The wave travel did not, to the extent of our observation, exhibit a predictable pattern, even when it recurred in the same region. This is exemplified in Figure 3E, where we traced the mean fluorescence intensity as a function of time in four adjacent ROIs, across approximately 50 µm in width from anterior to posterior. Over a 100 s interval, there were three large increases in the calcium signal (∼3-5 fold above the baseline), and the signal traveled in both proximal-distal and distal-proximal directions.

**Figure 3.**
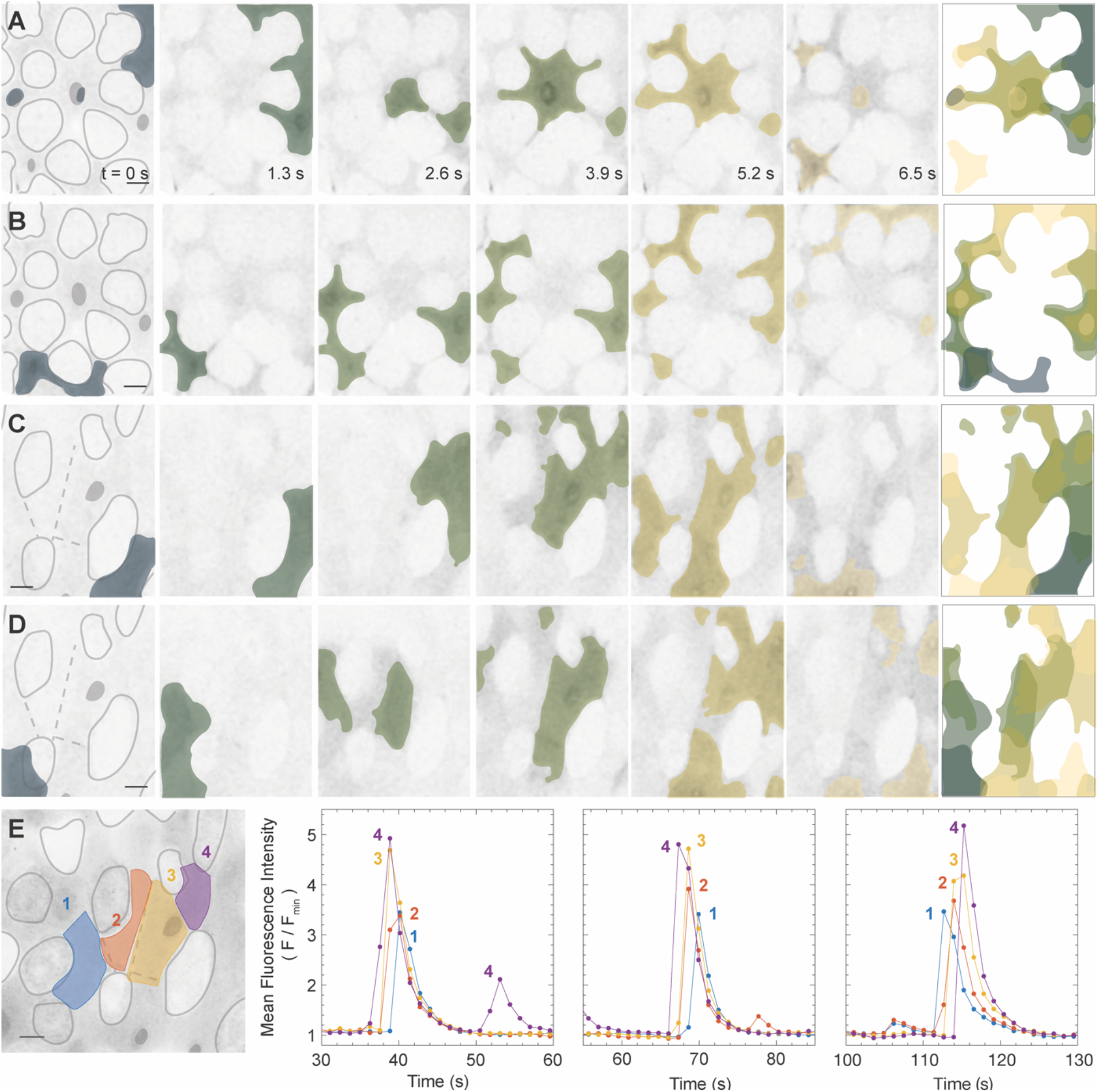
Calcium waves in interstitial cells (identified as the space between copper cells) exhibit a multitude of dynamic patterns. Calcium waves in two different fields from the same midgut are shown in A-B and C-D, demonstrating diverse spatial propagation dynamics in the same tissue. Approximations of copper cell outlines are depicted in gray. False coloring of the calcium signal is done using the scientific colormap bamako^32,33^. The signal transitions from dark to light color as a function of time with the last image illustrating a temporal maximum projection of the false colors. E) Four adjacent regions of interest (ROIs) with corresponding normalized mean fluorescence intensity of GCaMP as a function of time (*mex-GAL4>UAS-GCaMP*). Maximum intensity projections are shown. All scale bars, 10 µm. See Movie S1.

### Calcium oscillations in progenitor cells neither propagate from calcium waves in enterocytes nor correlate with oscillations in enteroendocrine cells

To directly test whether calcium oscillations in different cell types are coupled, we used orthogonal driver systems, *LexA/LexAop* ^35,36^ and *GAL4/UAS* ^26^, to express differently labelled calcium sensors in different cell types simultaneously, and performed multi-channel imaging to record the signal from both. From the StanEx collection of *LexA*-based enhancer trap drivers ^37^, we used the insertion *StanEx*^*SJH1*^, located upstream of the transcription start site for *escargot* to label progenitor cells. For convenience, we refer to this insertion as *esg-LexA* (Figure 4A-B, Figures S1 and S2). In combination with *mex-GAL4*, we can visualize calcium dynamics directly in enterocytes and progenitors, simultaneously (Figure 4C, D, Movie S4).

**Figure 4.**
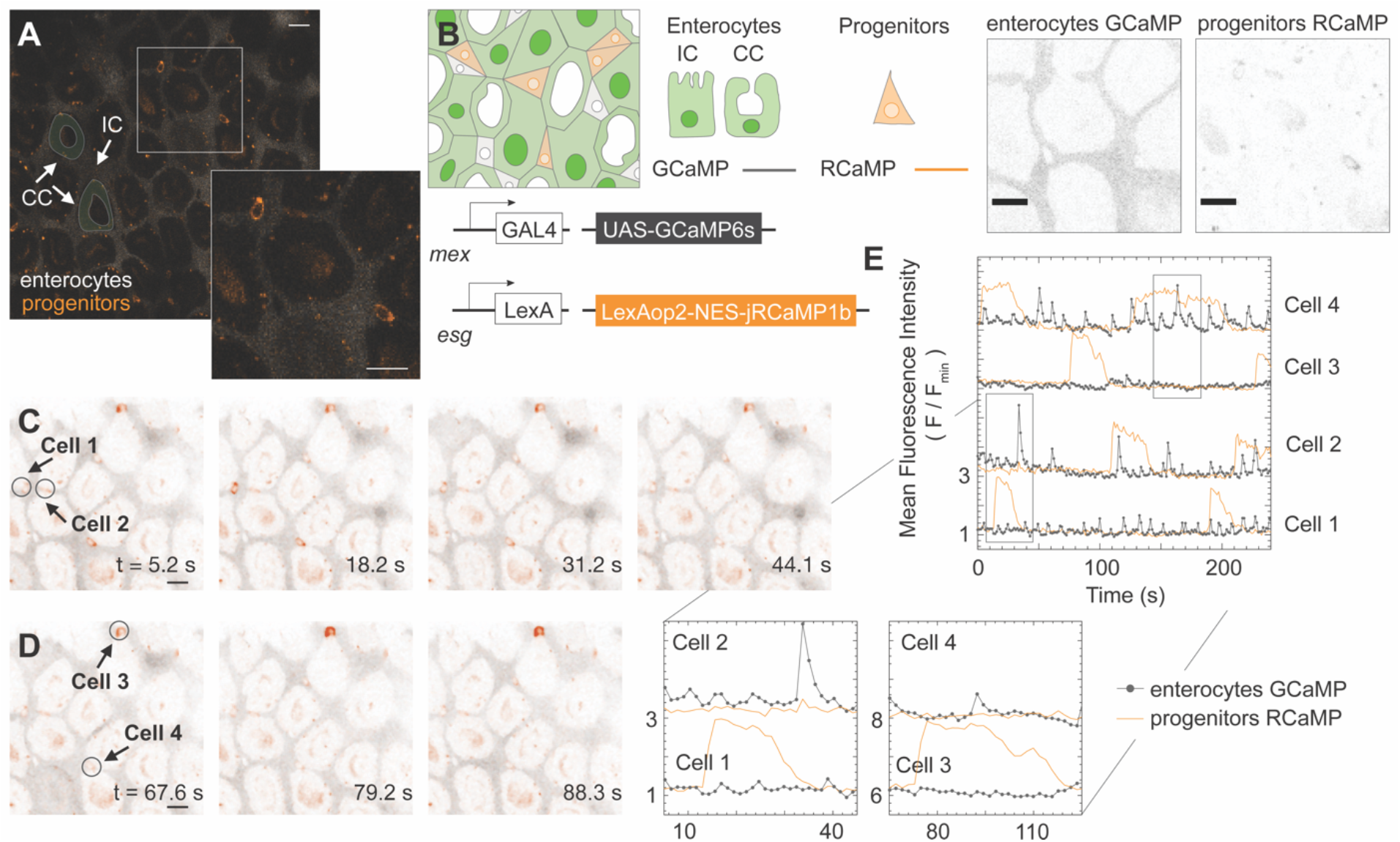
Calcium dynamics are independent between enterocytes and progenitors (Movie S4). A) Time point of a single-plane 240 s movie. Examples of enterocytes are marked with false-colored overlays and arrows. Cells are distinguishable for enterocytes (gray) and progenitors (orange pseudocolor). Inset shows a 2x magnification of the region within the square. B) Genetic design of the dual reporter line (*mex-GAL4>UAS-GCaMP, esg-LexA>LexAop-jRCaMP*). Cell types are distinguished by the expression of their fluorescent markers: enterocytes (GCaMP), and progenitors (RCaMP). C-D) Time lapse sequences from the single-plane movie showing calcium dynamics simultaneously in enterocytes and progenitors. An ROI was selected to encompass a single progenitor (‘Cell’) and to compensate for its movement throughout the movie. E) Mean fluorescence intensity of GCaMP (gray) and RCaMP (orange) as a function of time for the entire movie with insets that contain the time lapse sequences shown in C) and D) for the identified four progenitor cells. All scale bars, 10 µm.

ROIs were selected to cover a region a little larger than a single progenitor (Figure 1A), so that the cell of interest remained within the ROI even as the gut shifted due to the contracting muscle cells. Mean fluorescence intensities of the ROIs were traced as a function of time for both GCaMP (enterocytes) and RCaMP (progenitors) channels (Figure 4E). Figure 4E shows rapid oscillations, ∼24 mHz in the GCaMP channel and ∼20 s wide peaks in the RCaMP channel, characteristic of calcium dynamics in interstitial cells and progenitors, respectively (Figure 1D). If calcium waves propagated between enterocytes and progenitors, then the fluorescence intensity peaks should either overlap for the two channels or else one channel should consistently and immediately precede the other. We observed neither scenario. Instead, the signal increase in one channel was, as far as we could determine, independent of the other channel, and while there are situations where one signal precedes another, this occurrence was sporadic.

We also used the combination of the *GAL4/UAS* and *LexA/LexAop* systems to simultaneously visualize calcium dynamics in enteroendocrine cells (*pros-GAL4>UAS-GCaMP*) and progenitors (*esg-LexA>LexAOp-jRCaMP*). Similar to enterocytes and progenitors, we observed calcium oscillations in enteroendocrine cells and progenitor cells were independent of each other (Movie S5).

### Propagation of calcium waves and oscillatory calcium dynamics depend on functional gap junctions

Gap junctions are intercellular channels that allow direct transfer of small molecules and ions. Thus, cells sharing gap junctions are electrically coupled. For example, in the regenerative basal layer of the skin epithelium directed calcium signaling has been reported to be regulated by a major gap junction protein ^38^. Since the calcium waves appear to be travelling across several cell lengths, we hypothesized that calcium ions propagate across cells via gap junctions. To examine this hypothesis, we blocked gap junctions by adding the small-molecule inhibitor carbenoxolone ^17,39^ (CBX) to the imaging media for 15 min prior to to and during imaging of midguts with cell-type specific expression of calcium indicators. We analyzed the fluorescence intensity of calcium indicators in individual cells as a function of time for each cell type in control and CBX-treated guts (Figure 5D-E, Movies S6-8).

**Figure 5.**
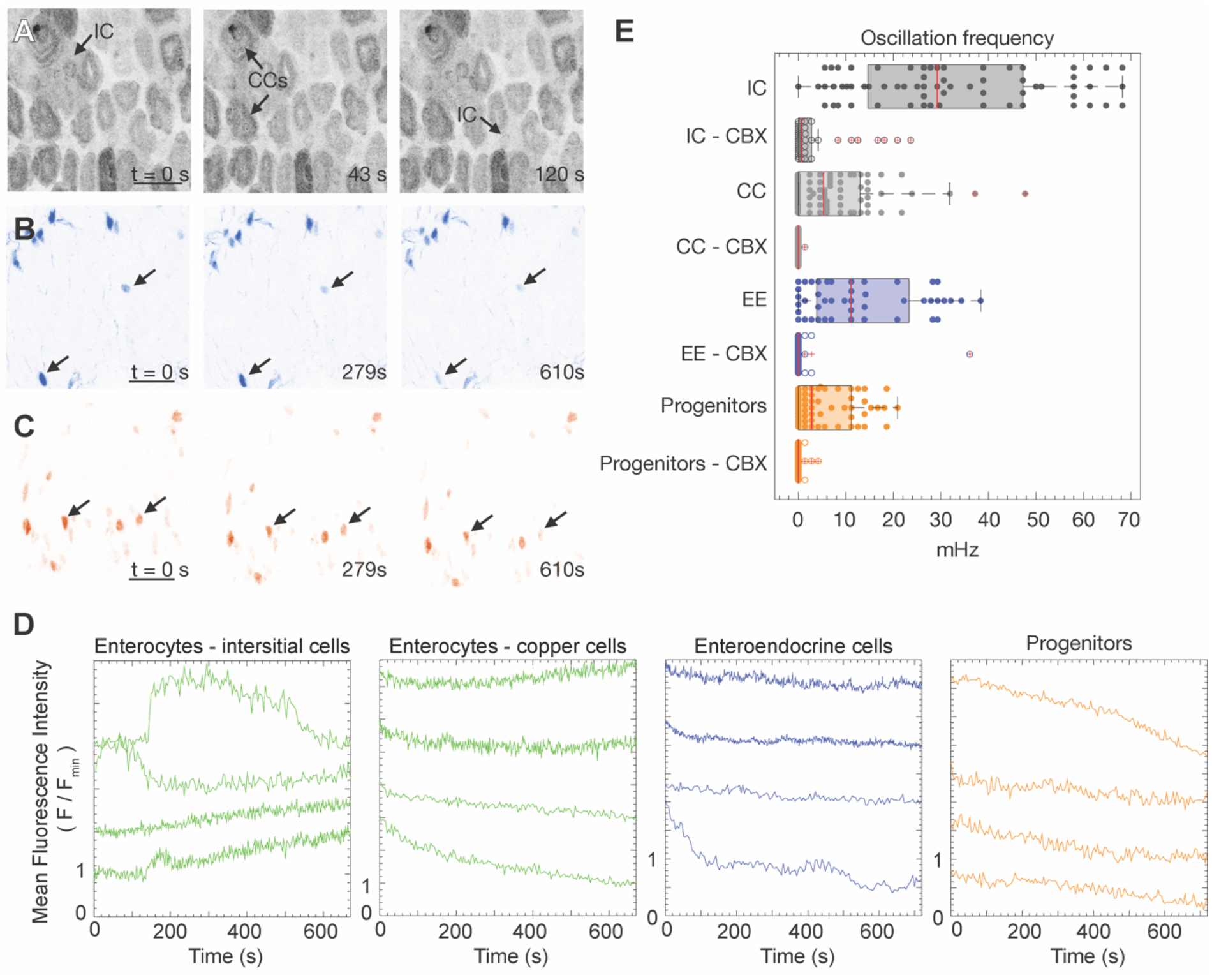
Propagation of calcium waves and oscillatory calcium dynamics depend on functional gap junctions. Time lapse image sequences of max intensity projections for tissue incubated with the gap junction inhibitor, carbenoxolone (CBX). A) Enterocytes (*mex-GAL4>UAS-GCaMP*, Movie S6) - interstitial cells (IC) and copper cells (CC), B) enteroendocrine cells (EE) (*pros-GAL4>UAS-GCaMP*, Movie S7), and C) progenitors (esg-GAL4>UAS-jRCaMP1b, Movie S8). All scale bars, 25 µm. D) Representative traces of fluorescence intensity of genetically encoded calcium indicators were selected from four movies for ECs, five movies for EEs, and three movies for progenitors. E) Comparison of calcium oscillation frequency for the different cell types in the middle midgut region with and without the gap junction inhibitor, carbenoxolone (CBX). The total number of ROIs analyzed per condition were IC: 65 (5 guts), IC - CBX: 38 (4 guts), CC: 74 (5 guts), CC - CBX: 44 ROIs (4 guts), EE: 45 ROIs (5 guts), EE - CBX: 51 ROIs (5 guts), progenitors: 59 ROIs (4 guts), progenitors - CBX: 39 ROIs (3 guts).

We found that gap junction inhibition eliminates calcium oscillations for all cell types. CBX treatment eliminated multicellular calcium waves that normally propagate through interstitial cells. CBX treatment also abrogated calcium oscillations in enteroendocrine cells (Figure 5B) and progenitors (Figure 5C). Intriguingly, CBX also caused calcium ions to accumulate in copper cells (Figure 5A); one possibility is that copper cells may act as a calcium sink when calcium flux through interstitial cells is blocked. Since the oscillations of progenitors are not coupled to enterocytes (Figure 4), this result was unexpected; potentially, whole-organ inhibition of gap junctions by CBX disrupts organ-scale calcium homeostasis with consequent inhibition of single-cell oscillations. Consistent with this notion, calcium oscillations in the intestinal stem cells have been shown to depend on both influx of calcium ions through the plasma membrane and on the internal release of calcium ions that have been actively sequestered in the intracellular stores by SERCA, Sarco-Endoplasmic Reticulum Calcium ATPase ^22^.

## DISCUSSION

Examining calcium dynamics in the midgut of adult *Drosophila*, we found that differently fated cells in the midgut R3 region each exhibit a characteristic pattern of calcium oscillations. Performing live imaging of midguts *ex vivo*, we identified propagation of calcium waves through networks of interstitial cells and characterized calcium oscillations in enteroendocrine cells and progenitors. Employing orthogonal expression of red and green calcium sensors, we demonstrated that the calcium dynamics of enterocytes and progenitors are paced independently of the other. We also found that the calcium dynamics of progenitors and enteroendocrine cells are also paced independently.

Our findings demonstrate that differentiated cell types in the same epithelium, despite deriving from the same progenitor cell population, exhibit calcium dynamics that differ from their mother cells and from each other. This result implies that as stem cell progeny undergo differentiation, they adopt fate-specific modes of calcium pacing. Calcium signal integration remains an intriguing open question due to its pleiotropic and yet ubiquitous nature, with one possibility that unique oscillatory calcium patterns are interpreted by downstream effectors to subsequently activate different cellular processes, and another that the calcium dynamics support the physiological functions of the specific cell types.

Intercellular calcium waves have been identified in several epithelial cell types in cell culture, and wave propagation, in many cases, is thought to occur via gap junctions ^40^. These cell-cell junctions are oligomers of connexins (innexins in *Drosophila*), with eight genes identified in *Drosophila* and encoding at least 10 transmembrane proteins ^41,42^. Transcriptomic analysis of the R3 region, indicates that enterocytes, enteroendocrine cells, and progenitors express predominantly Innexin 7, with trace expression of Innexin 2 ^43^. Our work demonstrated that whole-organ inhibition of gap junctions eliminated dynamic calcium responses in all three cell types and led to an accumulation of calcium ions in copper cells, and potentially disrupted organ-scale calcium homeostasis.

In summary, our findings establish a novel model for studying cell-type specific calcium dynamics within a heterogeneous population of stem and differentiated cells in an adult tissue. Our observations lay groundwork for using this highly tractable genetic model to investigate the roles of spatiotemporal calcium changes within signal transduction, organ renewal, and stem cell differentiation. Further studies need to examine the function of multicellular calcium waves in enterocytes and the specific roles of innexins in organ-scale calcium homeostasis.

## MATERIALS AND METHODS

### Fly stocks obtained from other sources

We obtained *20XUAS-IVS-NES-jRCaMP1b-p10* (BL63793), *20XUAS-IVS-GCaMP6s* (BL42746), *13XLexAop2-IVS-NES-jRCaMP1b-p10* (BL64428), and *P{ST*.*lexA::HG}SJH-1*^37^ (BL66632, referred to as *esg-LexA* in this paper, Figures S1 and S2) from the Bloomington Stock Center. *esg-GAL4* was obtained from the Kyoto Drosophila Genomics and Genetics Resource (DGRC). The following stocks were gifts: *mex-GAL4* (Carl Thummel), *esg-GFP[KI]/CyO* (Norbert Perrimon), and *pros-GAL4* (Sarah Siegrist). The complete key resources table can be found in Supplementary Information.

### Drosophila husbandry

Flies were fed a diet of standard cornmeal molasses food at 25°C or room temperature. Flies were collected 0-24 h post-eclosion, placed in vials with males and shifted to 25°C with 12 h light on and 12 h light off. The flies were fed a diet of standard cornmeal molasses food supplemented with yeast paste (Red Star, Active Dry Yeast), and the food vials were changed every 1-3 days. Experiments were performed on female flies, 4-7 days post-eclosion.

### Experimental setup

#### *Ex vivo* imaging

Female flies were briefly anaesthetized on carbon dioxide and then placed on ice in Eppendorf tubes to induce a chill coma. Guts were dissected in room temperature adult hemolymph-like (AHL) media (108 mM NaCl, 5 mM KCl, 2 mM CaCl_2_, 8.2 mM MgCl_2_, 4 mM NaHCO_3_, 1 mM NaH_2_PO_4_, 5 mM trehalose, 10 mM sucrose, 5 mM HEPES, pH 7.5, prepared by Electron Microscopy Sciences) and then bathed in whole organ *ex vivo* culture medium with 10 µg/ml isradipine (Selleck Chemical LLC) to decrease muscle contractions. The guts were then transferred in a droplet to #1.5 coverslips coated with poly-l-lysine (P4832-50ML, Sigma Aldrich) with 120 µm spacers (620001, Grace Bio-labs), and sealed with 250 µm thick PDMS sheets (see Figure S3). For experiments with the gap junction inhibitor, carbenoxolone was added to the *ex vivo* culture medium (100 µM CBX), and them tissue was incubated in this medium for 15 minutes prior to imaging.

#### Composition of whole organ *ex vivo* culture medium

Schneider’s Medium (21720024, Fisher Scientific) containing: 55 mM L-Glutamic acid monosodium salt (AAJ6342409 Alfa Aesar, Fisher Scientific, diluted from 1 M stock prepared in Schneider’s medium); 50 mM Trehalose (T5251-10G, Sigma Aldrich, diluted from 1 M stock prepared in Schneider’s medium)

2 mM N-Acetyl Cysteine (A9165-5G, Sigma Aldrich, diluted from a 200 mM stock prepared in sterile water)

1 mM Tri-sodium Citrate (PHR1416-1G, Sigma Aldrich, diluted from a 1 M stock prepared in Schneider’s medium)

5 mM HEPES (from 1 M solution, H0887-20ML, Sigma Aldrich)

#### Microscopy

An inverted Leica SP8 resonant scanning confocal microscope with a 40x/1.1 water-immersion objective was used to acquire movies that were analyzed in this study. Movies were captured at room temperature (20-23 °C). Confocal stacks were acquired with a z-step of 2 or 4 µm and typically contained 3-9 slices. For the complete list of movies and their information, including genotype, see Table S2. Movies were captured with cycle time of 4.7 s or faster. Movies were included in data analysis (Figure 5E) only if they were at least 290 s long. A Zeiss Z.1 Lightsheet was used with a 20x/1.0 objective by flowing a midgut into tubing (FEP009-031-B, Western Analytical) to obtain a high-resolution z-stack of the R3 region for Figure 1B.

#### Measurements of mean fluorescence intensity

Movie stacks were imported into Fiji ^44^ using the Bio Formats ^45^ plugin. In the acquired movie stacks, a plane was visually identified to be corresponding primarily to copper cells or to interstitial cells. In the selected plane, ROIs were drawn manually to correspond to the cell type of interest and mean fluorescence intensity values were obtained for each time point in Fiji. In the case of progenitors and enteroendocrine cells, the z-stacks were first converted into maximum projection movies in Fiji. Cells of interest were identified manually, and then tracked using the Active Contours ^46^ plugin in Icy ^47^ to record their mean fluorescence intensity. Mean fluorescence intensity values were plotted in relation to the minimum fluorescence values and peaks were identified using custom codes in Matlab 2019b (available upon request) with the aid of the Signal Processing Toolbox, and manually verified.

## ACKNOWLEDGEMENTS

The research was supported by the National Institutes of Health under grant R01 GM116000-01A1 (to L.E.O.) and R01 GM73164 (to D.J.M), the National Science Foundation CMMI 1834760 (to B.L.P.), and a seed grant from the Stanford Bio-X Interdisciplinary Initiatives Program (to L.E.O. and B.L.P.). A.A.K. was supported by the Swedish Research Council under the postdoctoral grant no. 2017-06156. M.M. was supported by NIH R01 GM124434. We acknowledge the use of the NRI-MCDB Microscopy Facility and the Resonant Scanning Confocal supported by the NSF MRI grant DBI-1625770, and the use of the Microfluidics Laboratory within the California NanoSystems Institute, supported by the University of California, Santa Barbara and the University of California, Office of the President. We thank the Bloomington *Drosophila* Stock Center (National Institutes of Health P40OD018537), the Kyoto Drosophila Genomics Resource Center Center (DGRC), and Carl Thummel, Norbert Perrimon, and Sarah Siegrist for fly stocks; and Jon-Michael Knapp for writing assistance.

