## Supplemental Information for "Independently paced calcium oscillations in progenitor and differentiated cells in an *ex vivo* epithelial organ"

### SUPPLEMENTARY INFORMATION

*Table S1 Key resources*

| Reagent type (species) or resource | Designation | Source or reference | Identifiers | Additional information |
|---|---|---|---|---|
| Genetic reagent (Drosophila melanogaster) | *UAS-jRCaMP1b* | Bloomington Drosophila Stock Center | BDSC: 63793; FLYB: FBti0180189 RRID: BDSC_63793 | *PBac{20XUAS-IVS-NES-jRCaMP1b-p10};* (Dana et al., 2016) |
| Genetic reagent (Drosophila melanogaster) | *UAS-GCaMP6s* | Bloomington Drosophila Stock Center | BDSC: 42746 FLYB: FBti0151344 RRID:BDSC_42746 | *P{20XUAS-IVS-GCaMP6s}* |
| Genetic reagent (Drosophila melanogaster) | *UAS-GCaMP6s* | | | *CyO/Sco; UAS-GCamp6s/TM6b,Tb*; shared by Craig Montell |
| Genetic reagent (Drosophila melanogaster) | *esgGAL4* | Kyoto DGRC | DGRC: 112304; FLYB: FBti0033872 | FlyBase symbol: *w[*]; P{w[+mW.hs]=GawB}NP0726/CyO* |
| Genetic reagent (Drosophila melanogaster) | *LeAop2-jRCaMP1b* | Bloomington Drosophila Stock Center | BDSC: 64428 FLYB: FBti0181971 RRID: BDSC_64428 | *P{13XLexAop2-IVS-NES-jRCaMP1b-p10}* |
| Genetic reagent (Drosophila melanogaster) | *lexA-esg* | Bloomington Drosophila Stock Center | BDSC: 66632 FLYB: FBti0185046 RRID: BDSC_66632 | *P{ST.lexA::HG};* (Kockel *et al.*, 2016) |
| Genetic reagent (Drosophila melanogaster) | *mexGAL4* | Shared by Carl Thummel | FLYB: FBgn0004228 | |
| Genetic reagent (Drosophila melanogaster) | *prosGAL4* | Shared by Sarah Siegrist | FLYB: FBgn0004595 | (Matsuzaki *et al.*, 1992) |

| | | | | |
|---|---|---|---|---|
| Genetic reagent (Drosophila melanogaster) | *GBE-Su(H)-GFP.nls* | PMID: 22522699 | | *w?; mw, GBE-Su(H)-GFPnls/(Cyo); Dr/TM6B* -- from (de Navascués et al., 2012) shared by Joaquin de Navascues |
| Genetic reagent (Drosophila melanogaster) | *UAS-his2b::CFP* | PMID: 28450412 | | *w; UAS-his2b::CFP/(Cyo);* + -- shared by Yoshihiro Inoue |
| Genetic reagent (Drosophila melanogaster) | *esg-GFP* | Shared by Norbert Perrimon | FLYB: FBrf0239545 | *esg-GFP[KI]/CyO* |
| Chemical compound | L-glutamic monosodium salt | Alfa Aesar | AAJ6342409 | 55 mM final concentration |
| Chemical compound | Trehalose | Sigma Aldrich | T5251-10G | 50 mM final concentration |
| Chemical compound | N-acetyl cysteine | Sigma Aldrich | A9165-5G | 2 mM final concentration |
| Chemical compound | Tri-sodium citrate | Sigma Aldrich | PHR1416-1G | 1 mM final concentration |
| Chemical compound | HEPES | Sigma Aldrich | H0887-20ML | 5 mM final concentration |
| Chemical compound, drug | Isradipine | Fisher Scientific, Selleck Chemical LLC | 50-153-5018; 50-136-1495 | 10 µg/m final concentration |
| Chemical compound, drug | Carbenoxolone | Sigma Aldrich | C4790 | 100 µM final concentration |
| Chemical compound | Poly-l-lysine | Sigma Aldrich | P4832-50ML | |
| Software, algorithm | Fiji | | RRID: SCR_002285 | Fiji (Schindelin et al., 2012), Bio Formats plugin |
| Software, algorithm | Icy | | | Icy (de Chaumont et al., 2012), Active Contours |
| Software, algorithm | Matlab | Mathworks | RRID: SCR_001622 | 2019b, Signal Processing Toolbox |

#### *Drosophila husbandry*

#### Fly stocks obtained from other sources

We obtained *20XUAS-IVS-NES-jRCaMP1b-p10* (BL63793), *20XUAS-IVS-GCaMP6s* (BL42746), *13XLexAop2-IVS-NES-jRCaMP1b-p10* (BL64428), and *P{ST.lexA::HG}SJH-1*(Kockel *et al.*, 2016) (BL66632, referred to as *esg-LexA* in this paper) from the Bloomington Stock Center. *esg-GAL4* was obtained from the Kyoto Drosophila Genomics and Genetics Resource (DGRC). The following stocks were gifts: *mex-GAL4* (Carl Thummel), *esg-GFP[KI]/CyO* (Norbert Perrimon), and *pros-GAL4* (Sarah Siegrist).

Flies were fed a diet of standard cornmeal molasses food at 25°C or room temperature. Flies were collected 0-24 h post-eclosion, placed in vials with males and shifted to 25°C with 12 h light on and 12 h light off. The flies were fed a diet of standard cornmeal molasses food supplemented with yeast paste (Red Star, Active Dry Yeast), and the food vials were changed every 1-3 days. Experiments were performed on female flies, 4-7 days post-eclosion.

**Comparison of *esg-LexA***

To understand the expression of *esg-LexA, esg-LexA>LexAop-jRCaMP1b* was co-expressed with *esg-GAL4>UAS-his2b::CFP* and *esg-GFP* in two separate experiments. Comparsion between e*sg-LexA>LexAop-jRCaMP1b* and *esg-GAL4>UAS-his2b::CFP* shows that *esg-LexA>LexAop-jRCaMP1b* almost always colocalizes with *esg-GAL4>UAS-his2b::CFP* (Figure S1). There are few cells that express *esg-GAL4>UAS-his2b::CFP*, but not *esg-LexA>LexAop-jRCaMP1b*, and this could be explained by that *LexAop-jRCaMP1b* is a transient signal. There are on occasion cells that express esg-*LexA>LexAop-jRCaMP1b* but not *esg-GAL4>UAS-his2b::CFP* (Figure S1 B). In general, *esg-LexA>LexAop-jRCaMP1b* exhibits a weaker signal.

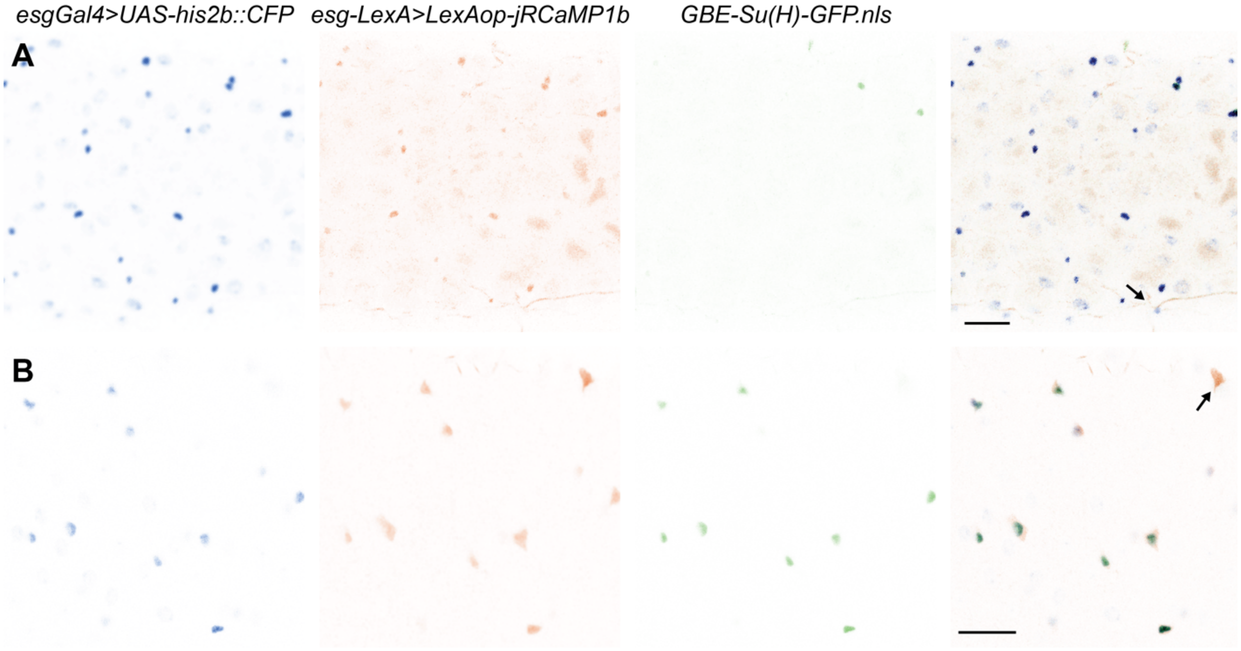


Figure S1 Co-expression of *esgGAL4>UAS-his2b::CFP* and *esg-LexA>LexAop-jRCaMP1b* in the A) middle midgut region and the B) posterior midgut. Few cells express *GBE-Su(H)-GFP* in the middle midgut compared to the posterior. The arrow in A) highlights an example of a cell that expresses *esgGAL4>UAS-his2b::CFP* but not *esg-LexA>LexAop-jRCaMP1b* and in B) the opposite. All scale bars, 25 μm.

*esg-LexA>LexAop-jRCaMP1b* also colocalized with *esg-GFP[KI]* (*esg-GFP*) and similarly, the signal is stronger in *esg-GFP*. A large majority of *esg-LexA>LexAop-jRCaMP1b* cells also express *esg-GFP*. There are several cells that express *esg-GFP* but not *esg-LexA>LexAop-jRCaMP1b*.

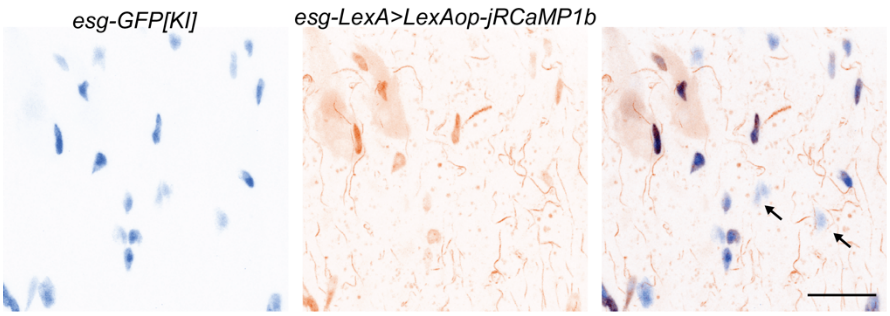


Figure S2 Co-expression of *esg-GFP* and *esg-LexA>LexAop-jRCaMP1b* in the copper cell region. The arrows highlight examples of cells that express *esg-GFP* but not *esg-LexA>LexAop-jRCaMP1b*. All scale bars, 25 µm.

### *Ex vivo imaging*

The assembled view of the *ex vivo* midgut mount for inverted microscopy is illustrated in Figure S3.

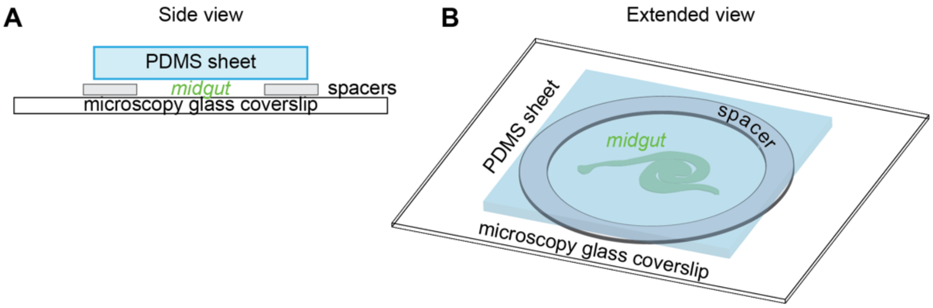


*Figure S3* Schematic of the *ex vivo* midgut mount for inverted microscopy. Spacer with a double-sided adhesive was attached to a microscopy coverslip coated with poly-l-lysine. The midgut was gently placed at the center in a drop of whole organ *ex vivo* culture medium. PDMS sheet was gently attached to the spacer. A) Side view and B) extended view.

### Additional files

Movie S1. Representative example of long-range calcium waves in enterocytes (interstitial cells, *mex-GAL4>UAS-GCaMP6s*) in a single plane. The movie was used to describe calcium waves in Figures 2 and 3. Scale bar is 25 µm. The movie is sped up 10X.

Movie S2. Maximal projection of calcium oscillations in enteroendocrine cells (*pros-GAL4>UAS-GCaMP6s*). Scale bar is 25 µm. Movie is sped up 100X.

Movie S3. Maximal projection of calcium oscillations in progenitors (*esg-GAL4>UAS-jRCaMP1b*). Scale bar is 25 µm. Movie is sped up 100X.

Movie S4. Maximal projection of calcium dynamics in enterocytes (*mex-GAL4>UAS-jRCaMP1b*) and progenitor cells (*esg-LexA>LexAop-jRCaMP1b*), simultaneously. Scale bar is 25 μm. Movie is sped up 10X.

Movie S5. Maximal projection of calcium dynamics in enteroendocrine (*pros-GAL4>UAS-GCaMP6s*) and progenitor cells (*esg-LexA>LexAop-jRCaMP1b*), simultaneously. Scale bar is 25 μm. Movie sped up 10X.

Movie S6. Maximal projection of enterocytes' response (*mex-GAL4>UAS-GCaMP6s*) to the gap junction inhibitor, carbenoxolone (100 μM). Scale bar is 25 μm. Movie is sped up 100X.

Movie S7. Maximal projection of enteroendocrine cells' (*pros-GAL4>UAS-GCaMP6s*) response to the gap junction inhibitor, carbenoxolone (100 μM). Scale bar is 25 μm. Movie is sped up 100X.

Movie S8. Maximal projection of progenitor cells' (*esg-GAL4>UAS-jRCaMP1b*) response to the gap junction inhibitor, carbenoxolone (100 μM). Scale bar is 25 μm. Movie is sped up 100X.

Table S2. List of movies analyzed and their related information, including genotype.

| Cell type | # z-slices | Spacing (µm) | Condition | Z-plane analyzed | Duration analyzed (s) | # of cells analyzed | Genotype |
|---|---|---|---|---|---|---|---|
| Copper cell (CCs) | 4 | 4 | - | 2 | 539 | 16 | *mexGAL4/esg-LexA>LexAop-jRCaMP1b; UAS-GCaMP6s* |
| | 6 | 4 | - | 4 | 719 | 11 | *mexGAL4/esg-LexA>LexAop-jRCaMP1b; UAS-GCaMP6s* |
| | 6 | 4 | - | 6 | 719 | 21 | *mexGAL4/esg-LexA>LexAop-jRCaMP1b; UAS-GCaMP6s* |
| | 6 | 4 | - | 1 | 293 | 15 | *mexGAL4/esg-LexA>LexAop-jRCaMP1b; UAS-GCaMP6s/Dr* |
| | 9 | 4 | - | 2 | 360 | 11 | *mexGAL4/esg-LexA>LexAop-jRCaMP1b; UAS-GCaMP6s/Dr* |
| | 5 | 4 | CBX | 2 | 719 | 10 | *mexGAL4/CyO; UAS-GCaMP6s* |
| | 6 | 4 | CBX | 1 | 719 | 12 | *mexGAL4/CyO; UAS-GCaMP6s* |
| | 6 | 4 | CBX | 4 | 719 | 12 | *mexGAL4/CyO; UAS-GCaMP6s* |
| | 6 | 4 | CBX | 1 | 1437 | 10 | *mexGAL4/CyO; UAS-GCaMP6s* |
| Interstitial cell (ICs) | 4 | 4 | - | 4 | 539 | 10 | *mexGAL4/esg-LexA>LexAop-jRCaMP1b; UAS-GCaMP6s* |
| | 6 | 4 | - | 5 | 719 | 12 | *mexGAL4/esg-LexA>LexAop-jRCaMP1b; UAS-GCaMP6s* |
| | 6 | 4 | - | 4 | 719 | 17 | *mexGAL4/esg-LexA>LexAop-jRCaMP1b; UAS-GCaMP6s* |
| | 6 | 4 | - | 4 | 293 | 15 | *mexGAL4/esg-LexA>LexAop-jRCaMP1b; UAS-GCaMP6s/Dr* |
| | 9 | 4 | - | 7 | 360 | 11 | *mexGAL4/esg-LexA>LexAop-jRCaMP1b; UAS-GCaMP6s/Dr* |
| | 5 | 4 | CBX | 2 | 719 | 9 | *mexGAL4/CyO; UAS-GCaMP6s* |
| | 6 | 4 | CBX | 2 | 719 | 12 | *mexGAL4/CyO; UAS-GCaMP6s* |
| | 6 | 4 | CBX | 2 | 719 | 7 | *mexGAL4/CyO; UAS-GCaMP6s* |
| | 6 | 4 | CBX | 2 | 1437 | 10 | *mexGAL4/CyO; UAS-GCaMP6s* |
| Entero-endocrine cell (EEs) | 5 | 2 | - | MAX | 720 | 8 | *UAS-GCaMP6s/CyO; prosGAL4* |
| | 4 | 4 | - | MAX | 495 | 6 | *UAS-GCaMP6s/CyO; prosGAL4* |
| | 6 | 4 | - | MAX | 716 | 7 | *UAS-GCaMP6s/esg-LexA>LexAop-jRCaMP1b; prosGAL4/Dr* |
| | 4 | 4 | - | MAX | 717 | 15 | *UAS-GCaMP6s/CyO; prosGAL4* |
| | 5 | 2 | - | MAX | 720 | 9 | *UAS-GCaMP6s/CyO; prosGAL4* |
| | 5 | 2 | CBX | MAX | 720 | 7 | *UAS-GCaMP6s/CyO; prosGAL4* |
| | 6 | 4 | CBX | MAX | 719 | 12 | *UAS-GCaMP6s/CyO; prosGAL4* |
| | 4 | 4 | CBX | MAX | 360 | 8 | *UAS-GCaMP6s/CyO; prosGAL4* |
| | 3 | 4 | CBX | MAX | 720 | 10 | *UAS-GCaMP6s/CyO; prosGAL4* |
| | 4 | 4 | CBX | MAX | 357 | 14 | *UAS-GCaMP6s/CyO; prosGAL4* |
| Progenitors | 6 | 4 | - | MAX | 430 | 8 | *esg-GAL4>UAS-his2b::CFP, GBE-Su(H)-GFP.nls/CyO; UAS-jRCaMP1b* |
| | 5 | 2 | - | MAX | 720 | 16 | *esg-GAL4>UAS-his2b::CFP, GBE-Su(H)-GFP.nls; UAS-jRCaMP1b* |
| | 4 | 4 | - | MAX | 717 | 14 | *esg-GAL4>UAS-his2b::CFP, GBE-Su(H)-GFP.nls/CyO; UAS-jRCaMP1b* |
| | 5 | 2 | - | MAX | 720 | 21 | *esg-GAL4>UAS-his2b::CFP, GBE-Su(H)-GFP.nls; UAS-jRCaMP1b* |
| | 6 | 4 | CBX | MAX | 719 | 13 | *esg-GAL4>UAS-his2b::CFP, GBE-Su(H)-GFP.nls/CyO; UAS-jRCaMP1b* |
| | 6 | 4 | CBX | MAX | 719 | 14 | *esg-GAL4>UAS-his2b::CFP, GBE-Su(H)-GFP.nls/CyO; UAS-jRCaMP1b* |
| | 6 | 4 | CBX | MAX | 719 | 12 | *esg-GAL4>UAS-his2b::CFP, GBE-Su(H)-GFP.nls; UAS-jRCaMP1b* |
